# Taphonomic damage obfuscates interpretation of the retroarticular region of the *Asteriornis* mandible

**DOI:** 10.1101/2023.06.13.544555

**Authors:** Abi Crane, Juan Benito, Albert Chen, Grace Musser, Christopher R. Torres, Julia A. Clarke, Stephan Lautenschlager, Daniel T. Ksepka, Daniel J. Field

## Abstract

*Asteriornis maastrichtensis*, from the latest Cretaceous of Belgium, is among the oldest known crown bird fossils, and its three-dimensionally preserved skull provides the most substantial insights into the cranial morphology of early crown birds to date. Phylogenetic analyses recovered *Asteriornis* as a total-group member of Galloanserae (the clade uniting Galliformes and Anseriformes. One important feature supporting this placement was enlargement of the retroarticular processes, which form elongate caudal extensions of the mandible in extant Galloanserae. Here, we reinterpret the jaw of *Asteriornis* and illustrate that the caudalmost portion of the mandibles are in fact not preserved. Instead, the caudal extremities of both the left and right mandibular rami extend to the surface of the fossil block containing the holotype skull, where they have eroded away. The originally identified retroarticular process of the right mandible—which exhibits a morphology and orientation strikingly similar to the retroarticular processes of certain extant and fossil galloanserans, including the early Palaeogene total-clade anseriforms *Conflicto* and *Nettapterornis—*instead represents a twisted and caudally displaced medial process. Nonetheless, anatomical comparisons with extant taxa reveal that we are unable to exclude the possibility that *Asteriornis* exhibited robust retroarticular processes comparable to those of extant Galloanserae. In light of the reinterpreted morphology of the *Asteriornis* mandible, we update the original anatomical character matrix used to investigate its phylogenetic relationships, and our revised phylogenetic analyses continue to support its position as a total-group galloanseran, as initially interpreted. We demonstrate additional morphological traits of the mandible supporting this phylogenetic position and provide new data on the nature and distribution of retroarticular processes among early crown birds.

## Introduction

*Asteriornis maastrichtensis* is one of only two well-represented Mesozoic taxa that have been confidently identified as members of Neornithes (crown group birds) (Clarke et al., 2005; Clarke et al., 2016; Field et al., 2020a; Mayr, 2022a), the sole clade of birds to survive the end-Cretaceous mass extinction (Longrich et al., 2011). The holotype and only known specimen, from the latest Cretaceous Belgium, consists of fragmentary post-cranial material and a nearly complete, three-dimensionally preserved skull (Field et al., 2020a). On the basis of phylogenetic analyses, *Asteriornis* was initially interpreted as a total-group galloanseran, exhibiting a unique combination of typically galliform and anseriform features suggesting a phylogenetic position close to the most recent common ancestor of Galliformes and Anseriformes. However, subsequent analyses including *Asteriornis* have instead recovered it as a stem-palaeognath (Torres et al., 2021; Musser and Clarke 2022), a member of the sister clade to the remainder of the avian crown group, and a notable result as it would constitute the only known representative of a Mesozoic total-clade palaeognath (Widrig et al. 2022), filling a major gap in the crown bird fossil record (Field et al. 2020b). The phylogenetic placement of *Asteriornis* has implications for interpreting the ancestral condition of crown group birds, and for informing divergence-time estimates for some of the deepest extant clades within the avian crown group. As such, a more complete and accurate understanding of the morphology and phylogenetic position of *Asteriornis* may provide important insight into the earliest stages of neornithine evolution.

Galliformes (landfowl) and Anseriformes (waterfowl) are now well understood to be sister groups occupying a position sister to the major extant clade Neoaves (Sibley & Ahlquist, 1990; Livezey, 1997; Mayr & Clarke, 2003; Ericson et al., 2006; Hackett et al., 2008; Mayr, 2008; Jarvis et al., 2014; Prum et al., 2015; Reddy et al., 2017; Kimball et al., 2019; Kuhl et al., 2021), but this relationship remained controversial prior to the widespread adoption of molecular phylogenetics (Olson & Feduccia, 1980; Ericson, 1996, 1997). First suggested as early as the 19^th^ century (Garrod, 1873), the clade Galloanserae (defined as the most recent common ancestor of Galliformes and Anseriformes, and all of its descendants (Mindell, 2020)) has hitherto only been supported by a limited number of morphological synapomorphies (Cracraft & Mindell, 1989; Ericson, 1996). Other previously proposed phylogenetic arrangements included Anseriformes as sister to the neoavian clades Charadriiformes (Olson & Feduccia, 1980; Feduccia, 1999) and ‘Ciconiiformes’ (Ericson, 1996), now recognised as a polyphyletic assemblage including storks, herons, ibises and flamingos (Ericson et al., 2006; Hackett et al., 2008; Jarvis et al., 2014; Kuramoto et al., 2015; Prum et al., 2015; Reddy et al., 2017; Kuhl et al., 2021), with Galliformes sometimes instead hypothesised to form a clade with Palaeognathae (Feduccia, 1999; Bourdon et al., 2010). Although the monophyly of crown Galloanserae is no longer controversial, the scarcity of osteological synapomorphies diagnosing the clade imposes challenges for identifying representatives of total-group Galloanserae in the fossil record.

The few osteological synapomorphies diagnosing Galloanserae are largely restricted to the skull (Ericson, 1996; Cracraft et al., 2001; Mayr, 2008; Bourdon et al., 2010; Mayr, 2011; Field et al., 2020a; Musser and Clarke 2022). Among these, the distinctive morphology of the pterygoid and the bicondylar quadrate-mandible articulation (Cracraft & Mindell, 1989; Ericson, 1996) have come under recent scrutiny, with analyses suggesting that they may instead represent retained symplesiomorphies of Neornithes rather than synapomorphies of Galloanserae (Mayr et al., 2018; Tambussi et al., 2019; Field et al., 2020a; Benito et al., 2022a). By contrast, the expanded, caudally projecting retroarticular process (which was considered convergent between galliforms and anseriforms in earlier work arguing against the monophyly of Galloanserae (Olson & Feduccia, 1980; Ericson, 1996, 1997; Olson, 1999), now stands as one of the few uncontroversial osteological synapomorphies of crown Galloanserae (Mayr, 2016; Field et al., 2020a), and this structure has played an important role in substantiating the assignment of several fossil birds to Galloanserae, including *Presbyornis* (Ericson, 1997; Livezey, 1997) and *Nettapterornis* (formerly *Anatalavis oxfordi*; Olson, 1999). Conversely, the apparent absence of enlarged, caudally projecting retroarticular processes has also been used to dispute putative galloanseran affinities for controversial fossil bird taxa, such as *Vegavis* (Mayr et al., 2018).

The retroarticular process is a caudally projecting process of the mandible extending caudal to the quadrate-mandible articulation, and its architecture typically comprises an extension of the angular bone within the mandibular postdentary complex (Baumel & Witmer, 1993; Vanden Berge & Zweers, 1993; Ericson, 1997). The retroarticular process acts as an area of insertion for the M. depressor mandibulae, and their size and shape are often correlated (Bock, 1964; Zusi, 1967; Ericson, 1997).

The retroarticular therefore plays an important functional role in the movement of the mandible, especially with respect to gaping and prying behaviour (Beecher, 1951; Zusi, 1967; Zweers & Berge, 1996; Previatto, 2012). Long retroarticular processes have been interpreted as adaptations for ecologies necessitating the beak to be opened against resistance (e.g., filter feeders; Bock, 1964; Zusi, 1967), or requiring a wide or forceful gape, as in some frugivorous or granivorous birds (Mayr, 2013). Among extant birds, well-developed retroarticular processes are uniformly present among representatives of Galloanserae, and are found in several neoavian taxa, including flamingos (Phoenicopteridae) and some representatives of Charadriiformes, Bucerotiformes, and Passeriformes (Beecher, 1951; Baumel & Witmer, 1993; Ericson, 1996; Zweers & Berge, 1996; Ericson, 1997; Mayr, 2005; Mayr et al., 2018; Field et al., 2020a; Mayr, 2022b). The morphology of the retroarticular process shows considerable variation among extant birds; the shape of the process ranges from being mediolaterally compressed and dorsoventrally deep in ducks (Anatidae) and flamingos, to a more rounded and mediolaterally wide form in landfowl (Galliformes) and screamers (Anhimidae) (Ericson, 1997).

Field et al. (2020a) interpreted *Asteriornis maastrichtensis* as possessing large and hooked retroarticular processes in their original description. The morphology of the *Asteriornis* retroarticular was considered comparable to that of the pan-anseriform *Nettapterornis oxfordi* (Olson, 1999), and was included as part of a suite of characters supporting galloanseran affinities for *Asteriornis* (Field et al. 2020a Fig. 1d). However, restudy of the CT data by Torres et al. (2021) resulted in a different interpretation, identifying the ‘retroarticular process’ of *Asteriornis* as a caudally displaced medial process of the mandible, twisted into a dorsoventral orientation. Further evaluation of high-contrast surface meshes confirms this observation and provides additional insight into the anatomy of the *Asteriornis* mandible. In its preserved position the process exhibits a dorsally directed hook and projects from the caudal extremity of the mandible, strongly resembling the retroarticular morphology of many anseriforms. Indeed, the overall shape of the retroarticular and medial processes in certain pan-galloanserans can be remarkably similar, as in the pan-anseriform *Conflicto* (Tambussi et al., 2019). When retrodeformed, however, this process shows a clear, although not exact, symmetry with the medial process of the left mandibular ramus.

**Figure 1.**
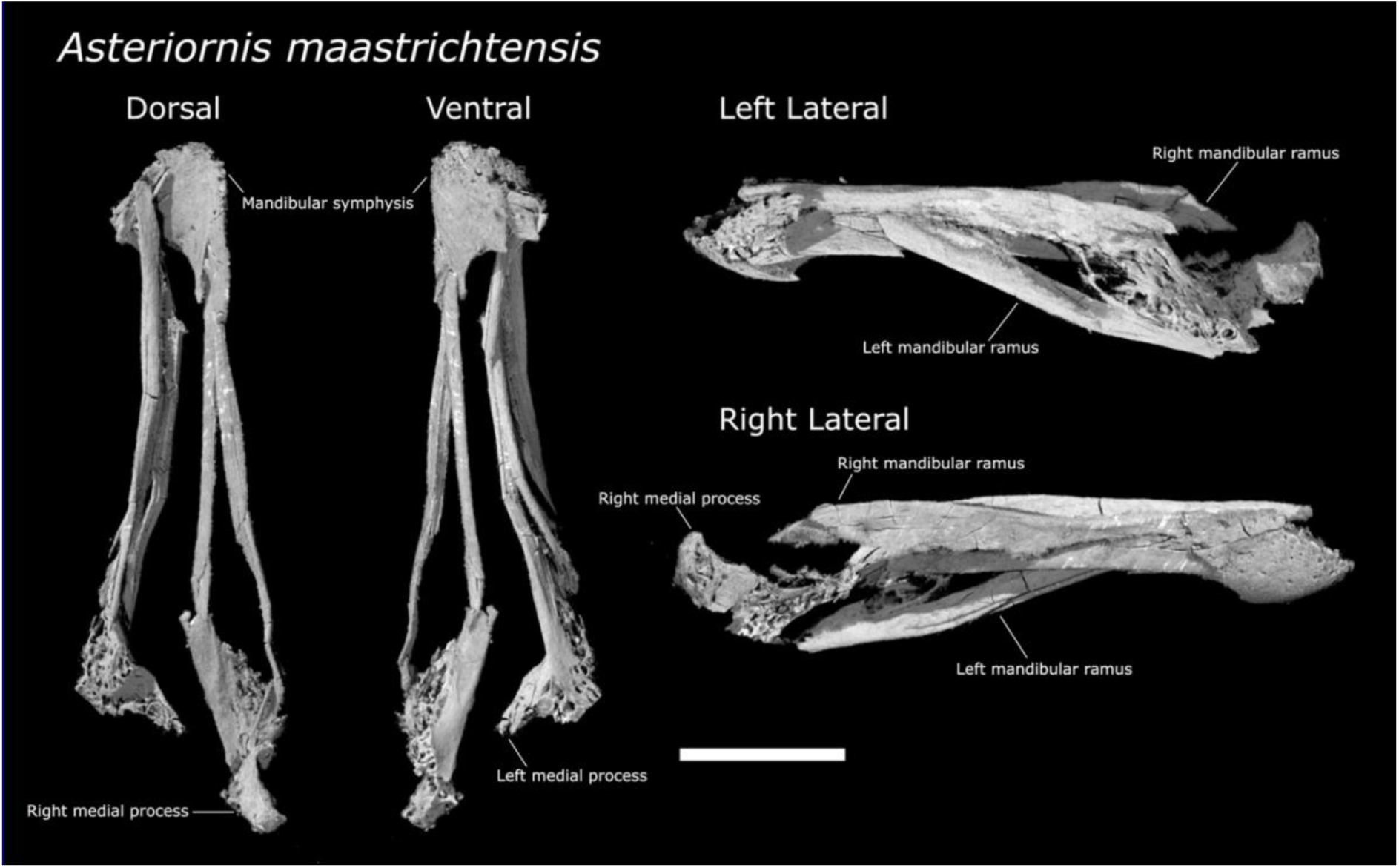
Mandible of *Asteriornis* holotype, as preserved. Whole mandible shown in dorsal, ventral, left and right lateral views. Scale bar equals 10mm.

Reinvestigation of the unprepared holotype of *Asteriornis* reveals that the caudal extremities of both the lower jaws eroded away at the surface of the fossil block (Fig. 4). We therefore sought to evaluate whether the state of preservation of the holotype mandibles could enable us to exclude the possibility that a galloanseran-like retroarticular was present in *Asteriornis*, which might have important implications for assessing the phylogenetic position of this crucial early neornithine, and could bear on our understanding of the evolutionary history of a key galloanseran synapomorphy. To assess the possibility of a retroarticular process having originally been present in *Asteriornis*, we compared the observable anatomy of the caudal ends of the preserved mandibles of *Asteriornis* with those of a range of extant bird taxa with their retroarticular processes intact and digitally removed. We also updated the phylogenetic matrix used by Field et al. (2020a) in light of our observations to assess their potential impact on the inferred phylogenetic position of *Asteriornis*.

## Methods

### Anatomical Comparisons

#### Taxon selection

Our sample included osteologically mature representatives of ten extant taxa. Six galloanserans were chosen to represent the spectrum of retroarticular morphologies represented within the clade. In this sample, retroarticular morphologies range from the mediolaterally narrow, ‘blade-like’ retroarticular diagnostic of Anseres (anseriforms other than Anhimidae) to the mediolaterally thicker processes typical of galliforms, which are more rounded in transverse section. We also included an aberrant specimen of *Meleagris gallopavo* with extremely elongated retroarticular processes atypical even for domestic members of this species in order to maximise the range of galloanseran retroarticular morphologies encompassed by our sample. Several non-galloanseran taxa were sampled for comparative purposes including two representatives of Palaeognathae (recovered as the closest extant relatives of *Asteriornis* by Torres et al. 2021) and two members of Neoaves: the flamingo *Phoenicopterus roseus*, which exhibits a large retroarticular process that may be convergently similar to that of anseriforms, and the shearwater *Puffinus puffinus*, which lacks a retroarticular process.

#### Imaging

All specimens were scanned using high-resolution computed tomography (CT); scanning was performed at the Cambridge Biotomography Centre (CBC) using a Nikon 49 Metrology XT H 225 ST high-resolution CT scanner. Scanned specimens were digitally segmented and rendered in VGSTUDIO MAX 3.4 (Volume Graphics, Heidelberg, Germany). In order to make anatomical comparisons between the broken caudal ends of the *Asteriornis* mandible and extant taxa, the caudal portions of the mandibles of the modern specimens (the ‘retroarticular regions’) were digitally removed. The retroarticular region was defined as including anything positioned caudal to the caudal end of the base of the medial process; this approximates the region of missing material from the mandibular rami of the *Asteriornis* holotype. This region was digitally removed using the clipping plane tool in VGSTUDIO MAX, and images were taken with and without the presence of this retroarticular region.

Due to an error during the original CT scanning process, the CT-data of *Asteriornis* studied by Field et al. (2020a) were left-right mirrored with respect to the actual fossil material; the scans used in the current work are corrected in reference to the original fossil. Features described as pertaining to the ‘left’ mandible by Field et al. (2020a) are therefore referred to as ‘right’ here, and vice versa.

### Phylogenetic analysis

We reanalysed the phylogenetic position of *Asteriornis* using a morphological dataset modified from Field et al. (2020a). We added one taxon to this dataset, the recently described putative stem-anseriform *Anachronornis*, based on scoring by Houde et al. (2023). For *Asteriornis*, we rescored characters 64 and 292 (Retroarticular process, shape), and 291 (Retroarticular process, presence) to unknown (see Results and Discussion). Additionally, we updated the scorings of *Vegavis* based on a recent redescription of the holotype by Acosta Hospitaleche and Worthy (2021). Finally, we rescored crown galliforms as having two sternal incisures (character 87) following comments on the homology of avian sternal trabeculae by Livezey and Zusi (2006), and rescored characters 271 and 272 as inapplicable for taxa lacking a hallux. The updated dataset consisting of 40 taxa and 297 characters was analysed under both maximum parsimony and Bayesian phylogenetic optimality criteria with molecular scaffolds derived from previous studies (Hackett et al., 2008; Gonzalez et al., 2009; Harris et al., 2014; Jarvis et al., 2014; Prum et al., 2015; Reddy et al., 2017; Kimball et al., 2019; Kimball et al., 2021; Kuhl et al., 2021; Simmons et al., 2022).

Maximum parsimony analyses were conducted in TNT v.1.5 (Goloboff & Catalano, 2016). After increasing the maximum number of trees to 99,999, a new technology search was run in which a minimum length tree was found in 10 replicates and default parameters were set for sectorial search, ratchet, tree drift, and tree fusion. After this, the maximum number of trees was set to 100 and a traditional search with default parameters was run on the trees in RAM to explore treespace more extensively. Absolute bootstrap frequencies were obtained from 1,000 replicates under a traditional search with default parameters.

Undated and tip-dated Bayesian phylogenetic analyses were conducted in MrBayes 3.2.7a (Ronquist et al., 2012) using the CIPRES Science Gateway (Miller et al., 2010), with tip-dating run under the fossilized birth-death model (Zhang et al., 2016). Priors and parameters followed those used by Field et al. (2020a), except as stated below. Age priors for extinct taxa were changed to uniform prior distributions spanning the age range of fossil occurrences, following recent work highlighting the importance of incorporating stratigraphic uncertainty into tip-dating analyses (Barido-Sottani et al., 2019; Püschel et al., 2020). Age ranges were based on previous literature (Benton & Donoghue, 2007; Mayr & Rubilar-Rogers, 2010; Ksepka & Clarke, 2015; Collinson et al., 2016; Worthy et al., 2017; Tambussi et al., 2019; Mayr et al., 2021; Houde et al., 2023), with updated ages for *Asteriornis* and *Vegavis* following recent stratigraphic studies of the Maastricht Formation and the Lopez de Bertodano Formation, respectively (Vellekoop et al., 2022; Roberts et al., 2023). Morphological synapomorphies were optimized under parsimony onto resulting tree topologies using TNT.

## Results and Discussion

### The retroarticular region of Asteriornis

The mandible of *Asteriornis* is taphonomically fragmented and the caudal ends of the mandibular rami are exposed at the surface of the block of the *Asteriornis* holotype (Fig. 1). The fossil is broken along a plane which has resulted in the loss of the caudal portion of the skull, including the occipital region, and the caudal extremities of the mandibles (Fig. 2a). The rostral and medial surfaces of the right articular region (Fig. 2c) are more completely preserved than the equivalent region of the left ramus, although this region of the mandible, including the medial process, has been taphonomically twisted, such that the long axis of the medial process is oriented dorsoventrally rather than mediolaterally. The tip of the medial process is well-preserved, showing a rostrally deflected tip, initially mistaken for a dorsally hooked tip as seen in the retroarticular processes of many anseriforms (Field et al. 2020a). The retroarticular process in the sampled extant taxa extends from the articular region on the caudal surface of the mandible in line with the long axis of the mandibular ramus (Fig. 3). Given the preservation of the right mandible of *Asteriornis*, it is clear that the region of the mandible that would have supported a retroarticular process, if one was present, is completely missing (Fig. 2c). The caudal portion of the right mandibular ramus is thus far too incomplete to exclude the possibility of a retroarticular process having originally been present.

**Figure 2.**
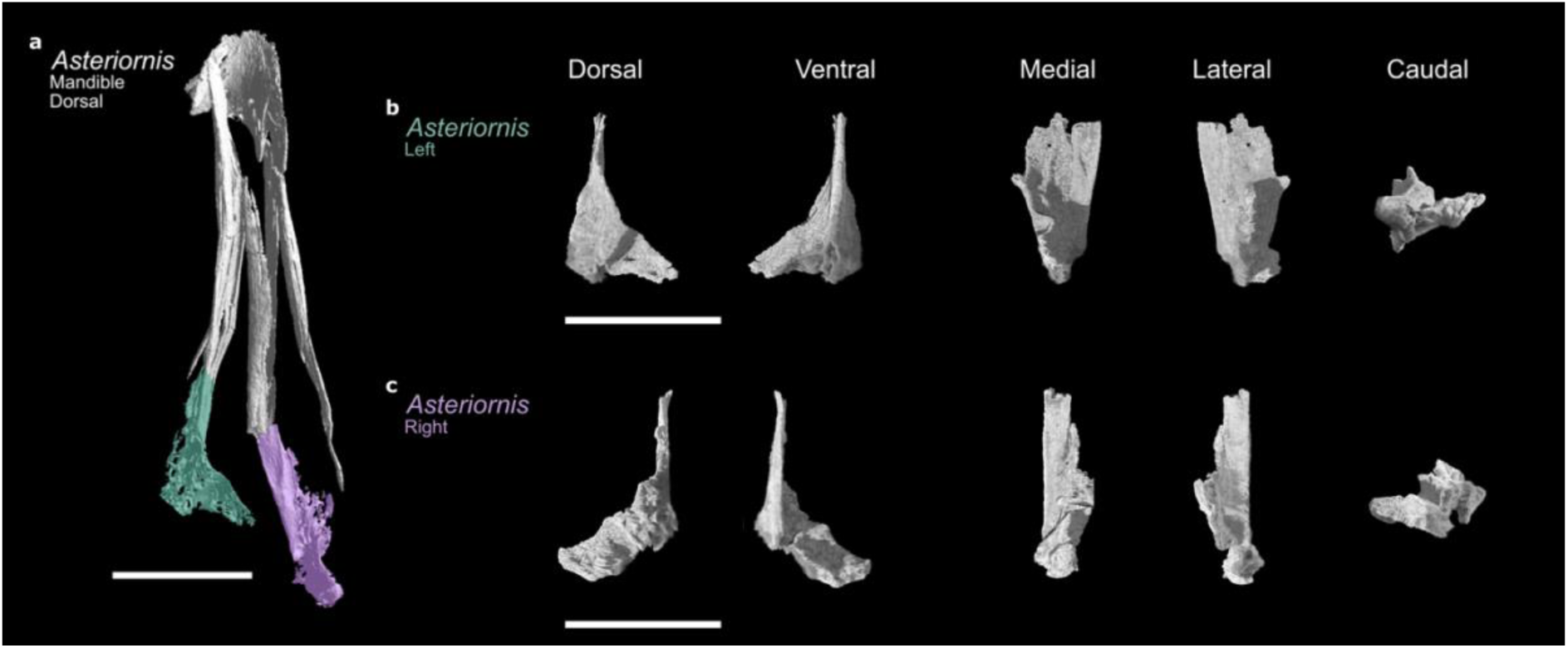
Caudal ends of the *Asteriornis* mandible. A) Whole mandible shown in dorsal view (with respect to mandibular symphysis). B,C) Caudal sections of right and left ramus shown separately in dorsal, ventral, medial, lateral and caudal views. B) Left ramus: dorsal surface left in medial view, right in lateral view. C) Right ramus: dorsal surface right in medial view, left in lateral view. Scale bar equals 10mm.

**Figure 3.**
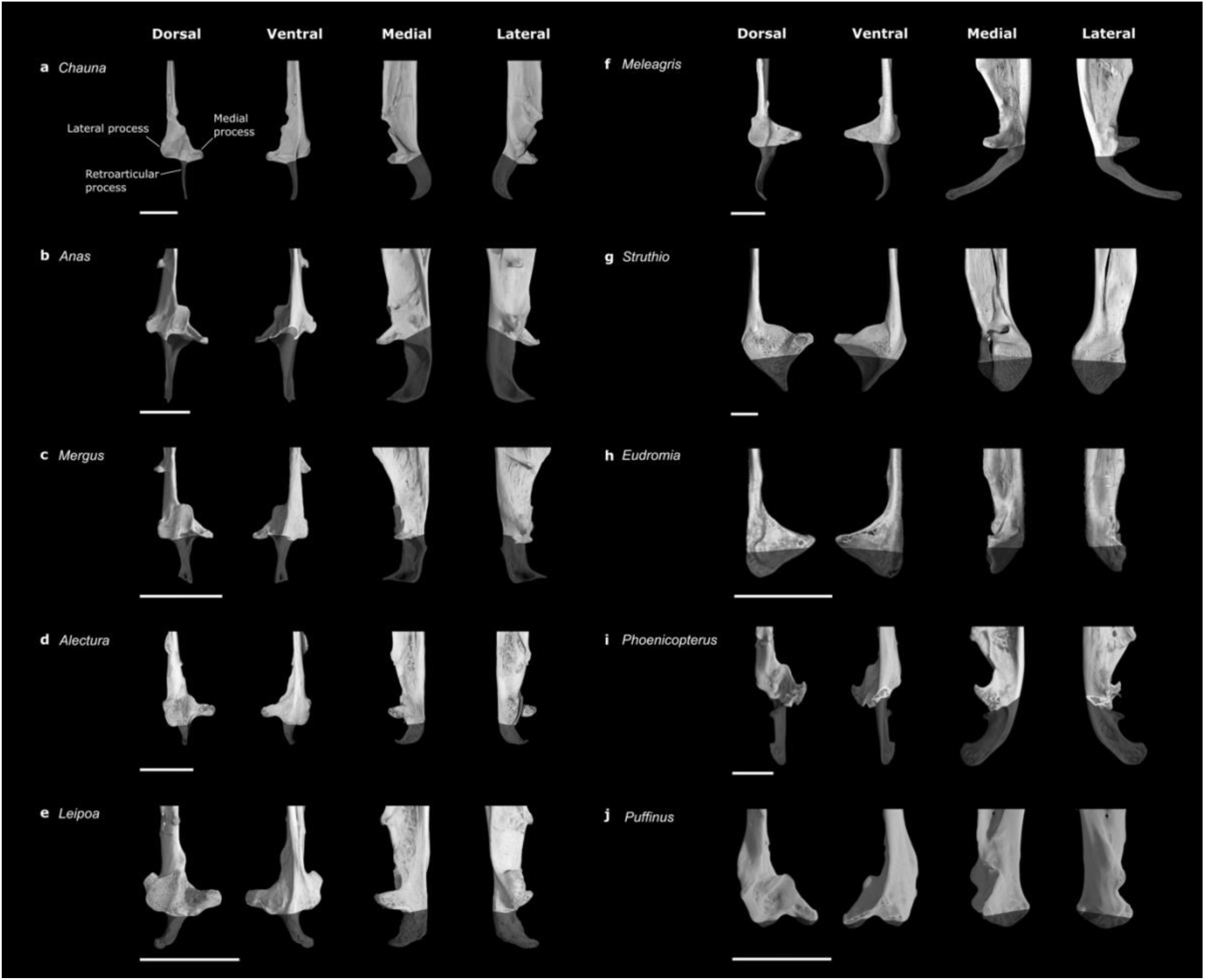
Caudal ends of the left mandibular ramus of a selection of extant taxa with retroarticular regions rendered transparent. Caudal ends of left mandibles in dorsal, ventral, medial and lateral views. Dorsal surface left in medial view and right in lateral view. Scale bar equals 10mm.

The left articular region is more fragmentary than the right articular region along its medial and rostral surfaces, yet is more complete than the right articular region along its lateral and caudal surfaces (Fig. 2b). The lateral process of the mandible is visible as a rostrocaudally elongate and mediolaterally shallow ridge extending along the lateral surface of the articular region. The caudal end of this process curves sharply towards the rostrocaudal midline of the mandibular ramus. Similarly, the caudal surface of the tip of the medial process appears to be preserved, and angles caudolaterally towards the middle of the mandibular ramus. The region between these two preserved caudal surfaces is broken, exposing the interior of the bone as an approximately triangular cross-section. The deepest part of this broken region is situated at the point where the break contacts the midline of the ramus towards the ventral surface, with the most caudally positioned intact portion of the mandible being the lateral process. As such, it appears that the original presence of a retroarticular process on the left mandibular ramus cannot be excluded either.

### Comparisons with extant taxa

The more complete caudal region of the left ramus of *Asteriornis* was compared with those of a range of extant taxa in order to determine whether the potential presence or absence of a retroarticular process in *Asteriornis* could be assessed (Fig. 3). The retroarticular processes of the surveyed taxa were digitally removed in order to approximate the preservation of this region in the *Asteriornis* holotype (Fig. 4). Given that the retroarticular process in extant taxa extends from the caudal surface in line with the long axis of the mandibular ramus (Fig. 3), comparisons with extant taxa illustrate that the preserved portions of the caudal surface of the articular region in *Asteriornis* do not include the region from which a potential retroarticular would extend (Fig. 4a), precluding any confident assessment of the original presence or absence of galloanseran-like retroarticular processes in *Asteriornis*.

**Figure 4.**
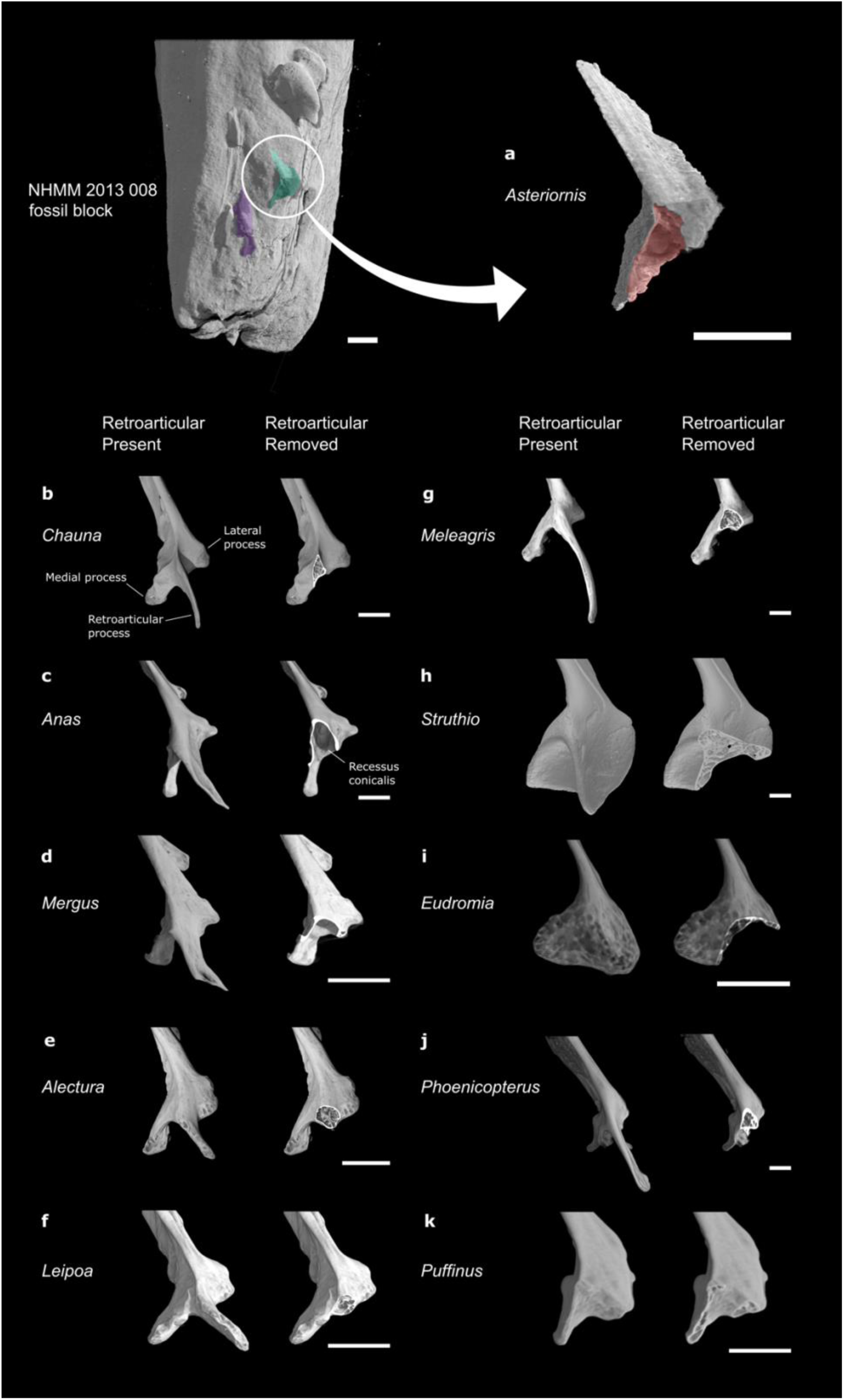
Caudal ends of the left mandibular ramus of *Asteriornis* and selected extant taxa. Extant taxa shown with retroarticular/caudal end of the mandible present and digitally removed. Positioned as viewed in the plane of exposure on the fossil block. Major region of broken/missing material of *Asteriornis* highlighted in red. Scale bar equals 5mm.

**Figure 5.**
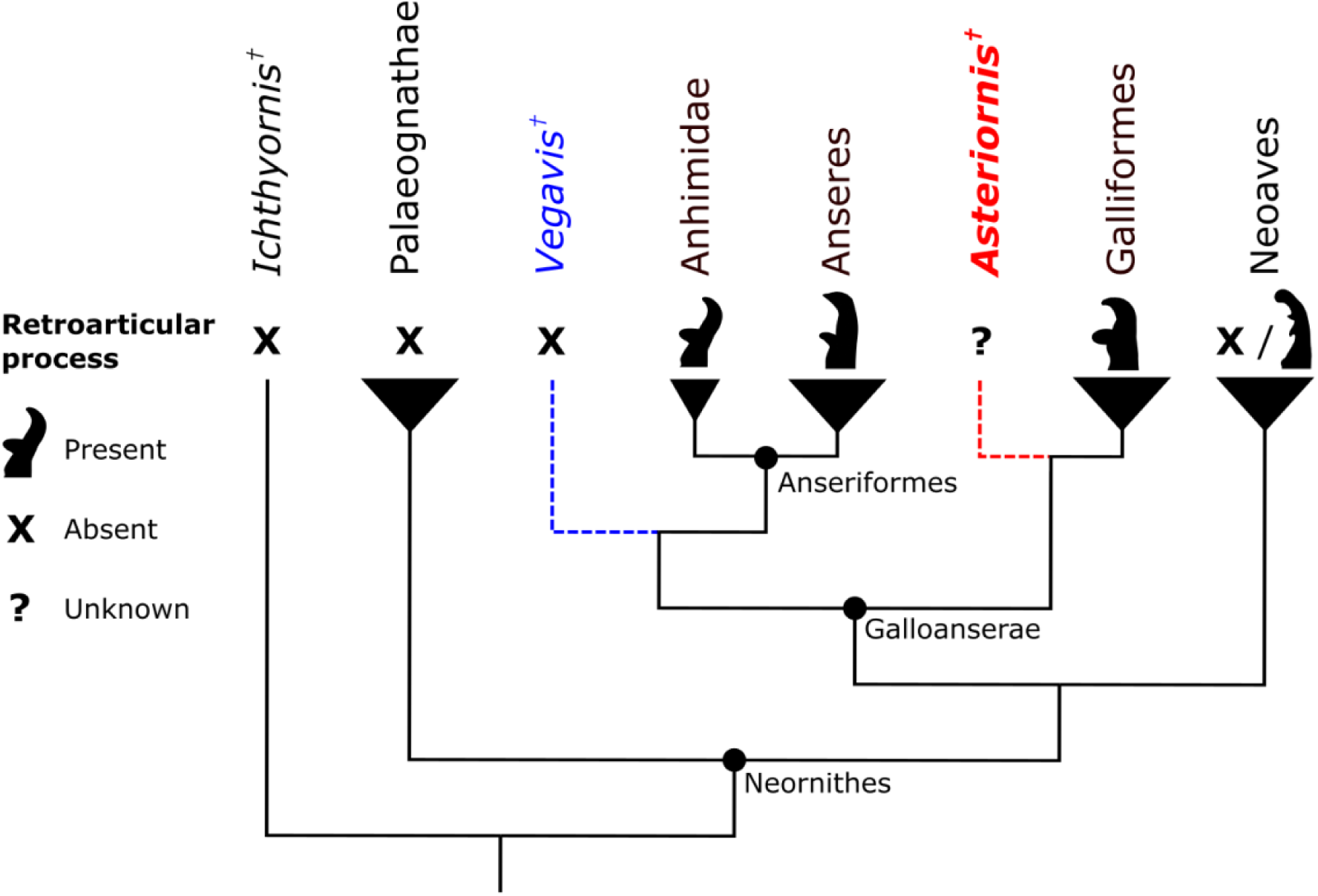
Cladogram showing recovered phylogenetic positions of *Asteriornis* and *Vegavis* under parsimony analyses. The presence, absence and uncertain presence of retroarticular processes is indicated, with silhouettes of a representative retroarticular morphology illustrated for those clades for which retroarticular processes are known.

### Implications for the phylogenetic position of *Asteriornis*

In the absence of evidence for enlarged retroarticular processes—an important galloanseran synapomorphy—the phylogenetic position of *Asteriornis* is in need of re-evaluation. Torres et al. (2021) reconstructed the mandible of *Asteriornis* with the medial process correctly positioned and identified in their supplementary information, although this was not discussed in that study. Their phylogenetic analysis found *Asteriornis* as a stem-palaeognath, albeit with weak statistical support. As such, the interpretation of *Asteriornis* as a stem-palaeognath was not emphasised in that study and was ascribed to a dearth of phylogenetically informative characters. The dataset used by Torres et al. (2021), an updated version of datasets used in Clarke (2004), Clarke et al. (2006), Huang et al. (2016), and Field et al. (2018) (though see Benito et al. (2022b) for critiques of this dataset), was primarily aimed at resolving stem-bird phylogeny with limited power to resolve neornithine phylogeny. Indeed, those authors reported that constraining *Asteriornis* to be a member of total-group Galloanserae in their analyses extended the length of their most parsimonious trees by only a single step. Moreover, no characters related to either the presence or morphology of the retroarticular process were included in the phylogenetic character matrix used in that study; thus, their reinterpretation of the originally identified retroarticular process of *Asteriornis* had no influence on its inferred phylogenetic position. Using an updated version of the same matrix, Benito et al. (2022a) found *Asteriornis* to be variably positioned as a stem-palaeognath under maximum parsimony, and as a crown anseriform under Bayesian inference, though the latter position is considered to be highly unlikely.

Musser and Clarke (2022) included *Asteriornis* in their new dataset for Galloanserae that includes more extant and extinct galloanserans than any previous dataset, with a particular focus on Anseriformes. *Asteriornis* was inferred as a stem-palaeognath in two analyses of that dataset: an analysis of morphological data under parsimony, and a Bayesian analysis of combined molecular and morphological data. However, that dataset contains a limited sample of palaeognaths, including only a tinamou and several lithornithids. In morphology-only analyses including the fossil anseriform *Wilaru tedfordi*, *Asteriornis* was inferred as a stem palaeognath with high bootstrap support. However, when *Wilaru* was excluded from their morphological analysis, *Asteriornis* was inferred as the sister taxon to Galloanserae with low support. Within both Bayesian analyses using combined data, *Asteriornis* was again recovered as a stem-palaeognath with low support.

Using the newly updated version of the Field et al. (2020a) dataset in which the presence and shape of the retroarticular were scored as unknown in *Asteriornis*, our maximum parsimony analysis recovered three most parsimonious trees (MPTs) with a consistency index (CI) of 0.255 and a retention index (RI) of 0.592. The strict consensus of all MPTs placed *Asteriornis* as a stem-galliform of unresolved affinities with respect to the Eocene taxon *Gallinuloides*, contrasting with the placement of *Asteriornis* on the galloanseran stem by the maximum parsimony analysis of Field et al. (2020a). As in Field et al. (2020a), statistical support for most deep divergences within the phylogeny was weak, with the clade uniting *Asteriornis*, *Gallinuloides*, and crown galliforms having a bootstrap support value of 26% and a Bremer support value of 2, likely due to the unstable positions of fossil taxa in the dataset. Bayesian analyses of the updated dataset were consistent with those of Field et al. (2020a) in recovering *Asteriornis* as stemward of *Gallinuloides* in tip-dating analysis, and as a stem-galliform crownward of *Gallinuloides* in undated analysis. Statistical support for placing *Asteriornis* crownward of *Gallinuloides* was low, with a posterior probability (PP) of 0.57, but support for an exclusive clade uniting *Asteriornis*, *Gallinuloides*, and crown galliforms was high (PP = 1 in undated analysis and 0.96 under tip-dating). These results support the initial interpretation of *Asteriornis* as a member of total-group Galloanserae, and potentially a member of total-group Galliformes (Field et al., 2020a).

Field et al. (2020a) reported that no retroarticular traits were inferred as synapomorphies of the *Asteriornis* + Galloanserae clade under maximum parsimony; as such, it is unlikely that the rescoring of characters related to the presence and morphology of the retroarticular process had a major effect on the inferred phylogenetic position of *Asteriornis* in that study. Therefore, the shift in the position of *Asteriornis* under maximum parsimony, from stem-galloanseran to stem-galliform, might instead be related to rescoring *Vegavis* and adding *Anachronornis*. Notably, the maximum parsimony and tip-dated Bayesian topologies we recovered were identical to those found by Houde et al. (2023), who also applied both of these changes to the Field et al. (2020a) dataset. As a further test of this hypothesis, we ran additional maximum parsimony analyses of our dataset in which scores for all retroarticular characters were reverted to those originally used by Field et al. (2020a). The resulting trees remained topologically unchanged from the results of our primary parsimony analysis (see Supplementary Material).

Although the presence of a retroarticular process cannot be confirmed in *Asteriornis*, additional characters support its placement as a total-group galloanseran and more broadly as a neognath. Under all of our recovered topologies, the presence of a rostromedial foramen in the otic process of the quadrate was optimized as a synapomorphy uniting *Asteriornis* with crown-group galliforms. This trait has also been documented in some other neognathous birds, including *Conflicto* and a few extant anseriforms, but not in palaeognaths (Tambussi et al., 2019; Field et al., 2020a).

*Asteriornis* additionally exhibits long, dorsally oriented medial processes of the mandible and a frontal depression forming a shallow, elongate groove, which are identified as synapomorphies of Galloanserae under some of our tree topologies. Furthermore, the presence of a subcapitular tubercle on the quadrate is widely found in Neognathae (Elzanowski & Stidham, 2010, 2011; Field et al., 2020a), and is supported as a potential synapomorphy of this clade by our analyses. Finally, we inferred two femoral characters observable in *Asteriornis* as possible synapomorphies of Neognathae: a caudally prominent medial supracondylar crest that interrupts the internal margin of the distal femoral shaft, as well as a poorly defined impression for the cranial cruciate ligament. These femoral characters are not found in any of the palaeognathous taxa sampled in our dataset, including Lithornithidae, a group of putative stem palaeognaths from the Paleogene. See Supplementary Material for further details of inferred synapomorphies from our analyses.

For comparison, we ran another maximum parsimony analysis in which *Asteriornis* was constrained to be a member of total-group Palaeognathae. Enforcing this constraint resulted in the recovery of two MPTs that were five steps longer than those found by our primary parsimony analysis. *Asteriornis* was recovered as the most stemward stem-palaeognath, whereas lithornithids were variably recovered as more crownward stem-palaeognaths or the sister group to *Tinamus*. Only a single unambiguous synapomorphy could be optimized in support for the clade uniting *Asteriornis* with crown palaeognaths: a splenial unfused to the dentary in adults, a character notably also present in extant megapodes (Galliformes: Megapodiidae). Of note, however, is the fact that the absolute age of *Asteriornis* at death remains unknown; thus, whether the unfused splenial of *Asteriornis* would have persisted in fully osteologically mature individuals cannot be unambiguously assessed. Overall, the results of our quantitative phylogenetic analyses and anatomical observations all suggest that a position within total-group Galloanserae is the best supported hypothesis for the affinities of *Asteriornis*, irrespective of our reinterpretation of its caudal mandibular anatomy.

### Morphology of the retroarticular region in crown birds

#### Galloanserae

The preserved caudal portion of the mandible of *Asteriornis* bears close resemblance to that of some galloanseran taxa after their retroarticular processes were digitally removed. Most notable are morphological similarities to the mandibles of the megapodes *Alectura lathami* and *Leipoa ocellata* (Fig. 4e,f), and the anhimid *Chauna torquata* (Fig. 4b), such as the shape and proportions of the lateral and medial processes. The lateral process of *Asteriornis* is most similar to that of *Alectura* and *Leipoa*, being relatively mediolaterally shallow and rostrocaudally long. The prominent medial process of *Asteriornis* is triangular in dorsal view and is similar in shape to that of *Chauna* and galliforms, whereas it is rostrocaudally narrower in anatids. Its moderate dorsal deflection in *Asteriornis* matches the condition in *Alectura*, *Leipoa* and *Chauna,* in contrast to the more steeply dorsally-pointed medial processes of anatids and other galliforms. Additionally, a sharp and prominent ridge along the ventral surface of the articular region in *Asteriornis*, in line with the long axis of the mandibular ramus, is similar to the condition in *Chauna*, but is developed to a lesser extent in *Alectura* and *Leipoa*. Importantly, Megapodiidae and Anhimidae represent the extant sister taxa to the rest of Galliformes and Anseriformes, respectively (Ericson, 1997; Livezey, 1997; Cracraft, 2001). Indeed, recent work suggests that other aspects of the jaw apparatus of megapodes and anhimids, namely the three-dimensional geometry of the quadrate, bear striking similarities to the inferred plesiomorphic condition of galloanserans (Kuo et al., 2023), corroborating earlier qualitative hypotheses (Elzanowski & Stidham, 2011; Elzanowski & Boles, 2012; Houde et al., 2023). The striking similarities between the preserved portions of the caudal mandible of *Asteriornis* and those of megapodes and anhimids is therefore in line with the hypothesis that *Asteriornis* may represent either a crownward stem galloanseran, or an early stem-galliform or stem-anseriform, as originally hypothesised by Field et al. (2020a).

Similarly, this region of the mandible of *Asteriornis* is similar to that of several fossil total-group galloanserans. Most notable is the resemblance to the caudal part of the mandible of the hypothesised early stem-anseriform *Conflicto* (Tambussi et al., 2019; Field et al., 2020a). The medial process is similarly proportioned in both taxa, and both show a slight rostral curvature at the tip, which is preserved most clearly on the right ramus of *Asteriornis*. The lateral process is also of similar size and shape in both taxa, presenting as a mediolaterally shallow, rostrocaudally oriented ridge along the lateral surface of the articular region. The prominence of the lateral processes and depth of the articular regions of the hypothesised total-group anseriforms *Nettapterornis*, *Anachronornis* and *Danielsavis* are also reminiscent of the condition in *Asteriornis* (Olson, 1999; Houde et al., 2023); the latter two taxa additionally exhibit a medial process with a shallowly dorsal orientation and a rostrally hooked tip similar to that of both *Conflicto* and *Asteriornis* (Houde et al., 2023).The morphological similarities in this region of the mandible of *Asteriornis* with those of fossil pan-anseriforms further support the interpretation of *Asteriornis* as a total-group galloanseran near the origin of the crown group.

By contrast, the caudal parts of the mandible of *Asteriornis* are markedly dissimilar to those of Anatidae, which also exhibit strikingly derived quadrates (Kuo et al. 2023). The anatids surveyed (*Anas platyrhynchos* and *Mergus albellus*; Fig. 4c,d) show a highly derived mandibular morphology including a large and deep depression (Ericson, 1997; Olson, 1999) between the mandibular ramus and the medial process, extending along the caudal surface of the latter, known as the recessus conicalis. The recessus conicalis is clearly exposed and visible upon removal of the retroarticular process in these taxa, and results in a distinctive morphology towards the rostral end of the articular region as the articular comprises part of this deep, caudally positioned depression (Fig. 4c,d). The regions of the caudal part of the mandible of *Asteriornis* which are preserved, including the medial process, clearly exclude the presence of a similarly deep, anatid-like recessus conicalis. The absence of this feature does not preclude stem-anseriform affinities for *Asteriornis*; although common to all members of Anatidae, a recessus conicalis is absent in Anhimidae and *Nettapterornis* (Olson, 1999) and is only shallow in *Anseranas* (the extant sister group to Anatidae) and in the fossil pan- anseriforms *Presbyornis* and *Conflicto* (Ericson, 1997; Tambussi et al., 2019). If an enlarged retroarticular process were present in *Asteriornis*, it may therefore be unlikely for it to have exhibited the extremely mediolaterally compressed, ‘blade-like’ morphology common to *Anseranas* and Anatidae (Fig. 3b,c), all of which also possess some form of recessus conicalis (Olson & Feduccia, 1980; Baumel & Witmer, 1993; Olson, 1999; Zelenkov & Stidham, 2018). We propose that the presence of a recessus conicalis and an anatid-like retroarticular process may be developmentally correlated, with the development of a recessus conicalis potentially constraining the mediolateral width of the retroarticular process by reducing the mediolateral extent of the caudal part of the mandible where the retroarticular attaches.

#### Extant non-galloanseran Neornithes

The caudal region of the mandibles of the palaeognaths *Struthio* and *Eudromia* (Fig. 4h,i) are morphologically dissimilar to that of *Asteriornis*. In both palaeognath taxa, the medial process is rostrocaudally broad and dorsoventrally shallow, especially in *Eudromia*, contrasting with the medial processes of *Asteriornis* and other extant galloanserans which are more pointed and dorsoventrally deep. The medial process of the palaeognaths is only very slightly dorsally deflected, as opposed to the shallow but obvious dorsal deflection of the process in *Asteriornis*. The lateral process is a small and indistinct dorsally-positioned ridge in *Struthio* and completely absent in *Eudromia,* contrasting with the distinct lateral process of *Asteriornis.* When the retroarticular region is digitally removed, the cross section of the remaining portion of the palaeognath caudal mandible is very different from the preserved caudal end of the *Asteriornis* mandible; the dorsal surface is prominently concave in both palaeognath taxa, while the exposed region is bulbous is *Struthio* (Fig. 4h) and uniformly thin dorsoventrally in *Eudromia* (Fig. 4i), in contrast to the broadly triangular cross-section of the articular region of *Asteriornis*. Even accounting for the uncertain presence of a retroarticular process, the morphology of the preserved parts of the caudal end of the mandible in *Asteriornis* are substantially more similar to those of galloanserans than those of palaeognaths. As such, the morphology of the caudal portion of the mandible casts further doubt on the total-group palaeognath affinities recovered for *Asteriornis* in some analyses (Torres et al., 2021; Benito et al., 2022b).

The preserved caudal end of the mandible of *Asteriornis* bears little morphological resemblance to those of the sampled neoavians, regardless of the presence of enlarged retroarticular processes. In *Puffinus puffinus* (Fig. 4k), which does not possess an enlarged retroarticular process, the medial process is angled so that the tip is directed caudally, extending to the same caudal position as the main body of the articular, and the caudal end of the mandible is much deeper dorsoventrally than that of *Asteriornis.* The lateral process is small and indistinct, presenting as only a slightly raised ridge on the lateral surface, and the medial process is proportionally small compared with that of the galloanserans examined. The dorsoventral depth of the articular region, the proportions of the medial and lateral processes, and the medially curved midline of the ramus at the caudal end yield a morphology of the caudal extremity of the mandible distinctly unlike that of *Asteriornis*. In *Phoenicopterus* (Fig. 4j), a neoavian with a retroarticular process that has been considered morphologically comparable to that of galloanserans (Ericson, 1996, 1997), the articular region is extremely deep dorsoventrally and narrow mediolaterally, with a medial process that is small and steeply angled dorsally. The medial process is pointed, with a sharp rostrally pointed hook, and the lateral process is completely absent. The retroarticular itself exhibits a unique morphology with a distinct dorsal process at its base and a slightly flared, rounded tip (Fig. 3i) unlike the sharp hooks seen in anatids (Fig. 3b,c).

#### Other fossil birds

The fact that the possible presence of an enlarged retroarticular process cannot be excluded in *Asteriornis* is particularly significant in the context of comparisons with other putative total-group galloanserans, notably *Vegavis* and Pelagornithidae. The mandible of *Vegavis* is known from a partial fragment of the postdentary complex, from which the absence of a large retroarticular has been inferred (Clarke et al., 2016; Mayr et al., 2018). Similarly, an enlarged retroarticular is known to be absent in pelagornithids (Bourdon, 2005; Mayr & Rubilar-Rogers, 2010). The galloanseran affinities of pelagornithids have recently been called into question; some characteristics thought to unite them with Galloanserae, notably a bicondylar quadrate-mandibular articulation and the morphology of the pterygoid, have instead been reinterpreted as possible crown bird plesiomorphies (Mayr et al., 2018; Tambussi et al., 2019; Benito et al., 2022b), and the morphology of the articular end of the pelagornithid mandible was considered to be very similar to that seen in the crownward stem bird clade Ichthyornithes (Mayr et al., 2021). By contrast, the lack of an enlarged retroarticular in non-neornithine members of Ornithurae such as *Ichthyornis* (Clarke, 2004) and Hesperornithes (Martin & Naples, 2008) supports the interpretation of this feature as a galloanseran synapomorphy. As discussed above, all extant galloanserans exhibit large and distinct retroarticular processes, as do known fossil pan-anseriforms such as *Anachronornis*, *Nettapterornis*, *Conflicto* and *Presbyornis* (Livezey, 1997; Olson, 1999; Tambussi et al., 2019; Houde et al., 2023), further suggesting that this morphology may have been present early in galloanseran evolutionary history. The confirmed absence of a retroarticular process in pelagornithids and *Vegavis*, in contrast to its unknown state in *Asteriornis*, is therefore worth considering in relation to the phylogenetic placement of these taxa and the distribution of this character through galloanseran evolutionary history.

## Conclusions

It is now understood that no retroarticular process is preserved in the holotype of *Asteriornis*; however, we cannot exclude the possibility of this feature having originally been present. The damage to the caudal ends of the mandibular rami of this specimen is extensive enough to have obscured any evidence that could unambiguously indicate the presence or absence of a retroarticular process. The portions of the mandible to which a possible retroarticular process would attach, based on comparisons with the mandibles of extant birds, are clearly missing from both mandibular rami, and thus it is currently impossible to exclude the possibility of a retroarticular process having originally been present in *Asteriornis*. This is in contrast to other putative total-group galloanserans for which the absence of a retroarticular process is known, including the only other well-established Mesozoic neornithine, *Vegavis* (Clarke et al., 2016), and the enigmatic extinct clade Pelagornithidae (Mayr & Rubilar-Rogers, 2010).

The position of *Asteriornis* within total-group Galloanserae is supported by our updated phylogenetic analyses reflecting the now unknown presence and morphology of the retroarticular process. Despite the uncertainty regarding this key galloanseran synapomorphy, other morphological features of the skull, and particularly the quadrate, continue to support galloanseran affinities for *Asteriornis*. This position appears to be further supported by novel observations of morphological similarities between the preserved caudal ends of the mandible of *Asteriornis* and those of modern galloanseran specimens where the retroarticular region was artificially removed. Most notable is the strong resemblance between the caudal part of the mandible of *Asteriornis* and the homologous region of megapodes and anhimids, the extant sister taxa to the rest of galliforms and anseriforms, respectively. This observation is consistent with the hypothesis of *Asteriornis* as a total-group galloanseran, phylogenetically proximal to the divergence between the galliform and anseriform lineages. Overall, this work revises our understanding of the mandibular morphology and phylogenetic position of *Asteriornis*, one of the oldest known neornithine birds and a critical datapoint in our evolving understanding of crown bird origins.

## Acknowledgements

We would like to thank M. Dickson for sharing details about the phylogenetic analyses run by Houde et al. (2023). J.B. and D.J.F. were funded by UKRI grant MR/S032177/1 to D.J.F. For the purpose of open access, the authors have applied a Creative Commons Attribution (CC-BY) licence to any Author Accepted Manuscript version arising.

